# Amplifying post-stimulation oscillatory dynamics by engaging synaptic plasticity with periodic stimulation: a modelling study

**DOI:** 10.1101/2024.01.25.577245

**Authors:** Jeremie Lefebvre, Aref Pariz

## Abstract

Periodic brain stimulation (PBS) techniques, either intracranial or non-invasive, electrical or magnetic, represent promising neuromodulatory tools for the treatment of neurological and neuropsychiatric disorders. Through the modulation of endogenous oscillations, PBS may engage synaptic plasticity, hopefully leading to persistent lasting effects. However, stabilizing such effects represents an important challenge: the interaction between induced electromagnetic fields and neural circuits may yield highly variable responses due to heterogeneous neuronal and synaptic biophysical properties, limiting PBS clinical potential. In this study, we explored the conditions on which PBS leads to amplified post-stimulation oscillatory power, persisting once stimulation has been turned off. We specifically examined the effects of heterogeneity in neuron time scales on post-stimulation dynamics in a population of balanced leaky-integrated and fire (LIF) neurons that exhibit synchronous-irregular spiking activity. Our analysis reveals that such heterogeneity enables PBS to engage synaptic plasticity, amplifying post-stimulation power. Our results show that such post-stimulation aftereffects result from selective frequency- and cell-type-specific synaptic modifications. We evaluated the relative importance of stimulation-induced plasticity amongst and between excitatory and inhibitory populations. Our results indicate that heterogeneity in neurons’ time scales and synaptic plasticity are both essential for stimulation to support post-stimulation aftereffects, notably to amplify the power of endogenous rhythms.

## Introduction

Brain stimulation has attracted significant interest in the last decades [1–3]. Various simulation techniques have shown promising results, and more are coming. Researchers, experimentally and theoretically, have addressed the numerous challenges related to the effects of these interventions on behaviour [4,5], brain function [6], as well as pathologies such as epilepsy [1,7], Parkinson [8], major depressive disorder (MDD) [9, 10] and stroke [11, 12]. Despite these promising results, it is still unclear how brain stimulation interventions shape endogenous brain dynamics [13–16] and the neural circuits that support them [17,18]. Indeed, brain stimulation outcomes remain variable: induced changes in neuron ‘s excitability vary remarkably between stimulation sites, repeated trials, and subjects, oftentimes vanishing after stimulation offset [19–22]. Uncovering the source of this variability can help to optimize existing brain stimulation paradigms and stabilize their effect on brain dynamics and plasticity.

Periodic brain stimulation (PBS) techniques, such as transcranial alternating current stimulation (tACS) and repetitive transcranial magnetic stimulation (rTMS), have repeatedly been shown to be capable of altering neurons’ dynamics to interfere with cortical rhythms [14, 15, 23–28]. The proposed mechanism of action of these techniques is the use of periodic electromagnetic waveforms to engage endogenous oscillatory activity to provoke structural and functional changes within targeted regions [27, 29]. Time-varying currents have indeed been shown to entrain endogenous oscillations [15,16] to support persistent effects that outlast stimulation duration [15, 30]. Induced over-lasting effects, such as increase and/or decrease in spectral power, may last for several minutes to hours after stimulation offset [17, 24, 25]. Multiple hypotheses for such persistent effects have been proposed, ranging from feedback reverberation [15, 31] to synaptic plasticity [19, 32]. Yet, mechanisms remain poorly understood and outcomes highly variable [33–35]. The efficacy of entrainment and subsequent post-stimulation effects are state-dependent [15,36], notably because of the competing influences of endogenous oscillations and PBS [30, 36].

The physiological changes involved in mediating post-stimulation after-effects, which are believed to support the reported clinical efficacy of PBS interventions, warrant further investigation. Most computational models fail at capturing these over-lasting effects. One central challenge stems from biophysical heterogeneity. Indeed, the influence of time-varying PBS on neural activity and plasticity relies heavily on biophysical cellular properties, such as the membrane time constant (MTC) [18]. The MTC is a quantity that reflects the agility of neurons in response to time-varying stimuli [37], and dictates their varied frequency selectivity [18]. The MTC varies significantly across cortical layers, and brain areas, ranging from a few to tens of milliseconds [38, 39]. Such variability has been shown to mediate selective, direction-specific synaptic plasticity under tACS [18] and hence represents a promising candidate in supporting persistent post-stimulation effects. Indeed, stimulation-induced and MTC-dependent changes in neuronal spike timing, further modulated by endogenous oscillations, may solicit Hebbian spike-timing dependant plasticity (STDP) to support lasting changes in synaptic weights [17, 18].

To better understand the mechanisms mediating post-stimulation aftereffects, one may first examine the effect of periodic stimulation on synchronous neuronal populations to determine how entrainment engages STDP synaptic plasticity in the presence of MTC heterogeneity [17,18,25]. In this study we explore the circumstances under which the application of electrical stimulation induces post-stimulation aftereffects in the form of an enhancement in endogenous oscillation power [34,40,41]. We characterized the relationship between neurons’ MTC variability, stimulation parameters (i.e., amplitude, frequency), and how they collectively shape synaptic plasticity amongst different groups of neurons as well as on the power of endogenous rhythmic activity. The variable responses of neurons, resulting from differences in MTCs, were found to be essential in supporting the amplification in endogenous oscillatory power [17, 24, 25]. The effect was further found to be stimulation amplitude- and frequency-specific. Our results reveal that particular synaptic connections, namely between excitatory and excitatory (E→E) and between inhibitory and excitatory (I→ E), are centrally involved in generating post-stimulation oscillatory amplification. The results suggest considering the effects of heterogeneity in biophysical attributes in designing and optimizing brain stimulation intervention protocols.

## Results

Besides entrainment, which naturally occurs through the oscillatory modulation of targeted regions [27], one purpose of PBS is to yield persistent effects that outlast stimulation duration. Intuitively, this objective can not be fulfilled unless PBS changes some physiological characteristics of the area under intervention. This can manifest itself through changes in the connectivity, cellular features, or both [42–44]. While sufficiently large amplitude stimulation is capable of altering neuronal spiking activity [18], the nature of the responses will depend on the neurons’ heterogeneous biophysical attributes. Such heterogeneity challenges our ability to reconcile the effects of PBS on neuronal dynamics, synaptic plasticity and possible over-lasting effects, which are a central component of PBS clinical efficacy. Such a key attribute is the membrane time constant (MTC). The membrane time constant is a key parameter representing the agility of neurons in response to time-varying stimuli [18, 37, 45]. However, MTC varies across neuron types, brain regions, and cortical layers [18, 38, 39]. Such wide heterogeneity in time scales translates into significant variability in neurons’ response to periodic stimulation: neuron spiking phase depends on the interplay between stimulation frequency and the neurons’ MTCs [18]. For instance, in the leaky integrate and fire (LIF) neuron model used in this study (see *Materials and methods*), differences in the spiking phase (Δ*ϕ*(*τ*_*m*_)) resulting from a stimulation frequency *ω*_*s*_ between neurons with distinct MTCs can be translated into a difference in spike timing i.e. Δ*T* = Δ*ϕ*(*τ*_*m*_)*/ω*_*s*_. Such difference in spike timing [46] has important implications for Hebbian plasticity, stimulation-induced changes in synaptic weights, and their joint influence on endogenous oscillatory activity.

Previous studies have shown that heterogeneity in neuronal timescales promotes selective, direction-specific synaptic modifications [18]. This occurs because heterogeneity establishes a temporal hierarchy between neurons with different MTCs, promoting the potentiating and/or depression of synapses, something one can capitalize upon using PBS. This selectivity was found to play an important role in mediating persistent PBS oscillatory aftereffects. We built a network of 10000 (8000 excitatory (E) and 2000 inhibitory (I)) LIF neurons with 10% of connection probability with plastic synapses representing a cortical network (see *Materials and methods* section and table 1). With these parameters, the network is found in a Synchronous-Irregular (SI) balanced state [48], with a peak frequency within the *β* rhythm (*f* = 28±1 *Hz*). Note that throughout this manuscript we use *f* to refer to the endogenous frequency (i.e., the frequency observed in the network without stimulation), and *ω*_*s*_ refers to the exogenous frequency (i.e., the stimulation frequency). To promote entrainment and increase the signal-to-noise ratio (i.e. endogenous oscillations vs. PBS) we set our system in a weak coupling regime. Having this condition helps avoid the competition between recurrent synaptic input and stimulation-induced fluctuations [30, 36]. Note that although individual synaptic weights might appear small, the net amplitude of synaptic currents is comparable to the stimulation-induced fluctuations in membrane potential: the average of maximum neurons’ synaptic current, and its standard deviation is ∼ 0.5*mV* and∼ 0.1, respectively. To model plasticity, we employed Hebbian spike-timing dependent plasticity (STDP) amongst *E*→ *E, E*→ *I*, and *I*→ *E* neurons.

**Table 1.**
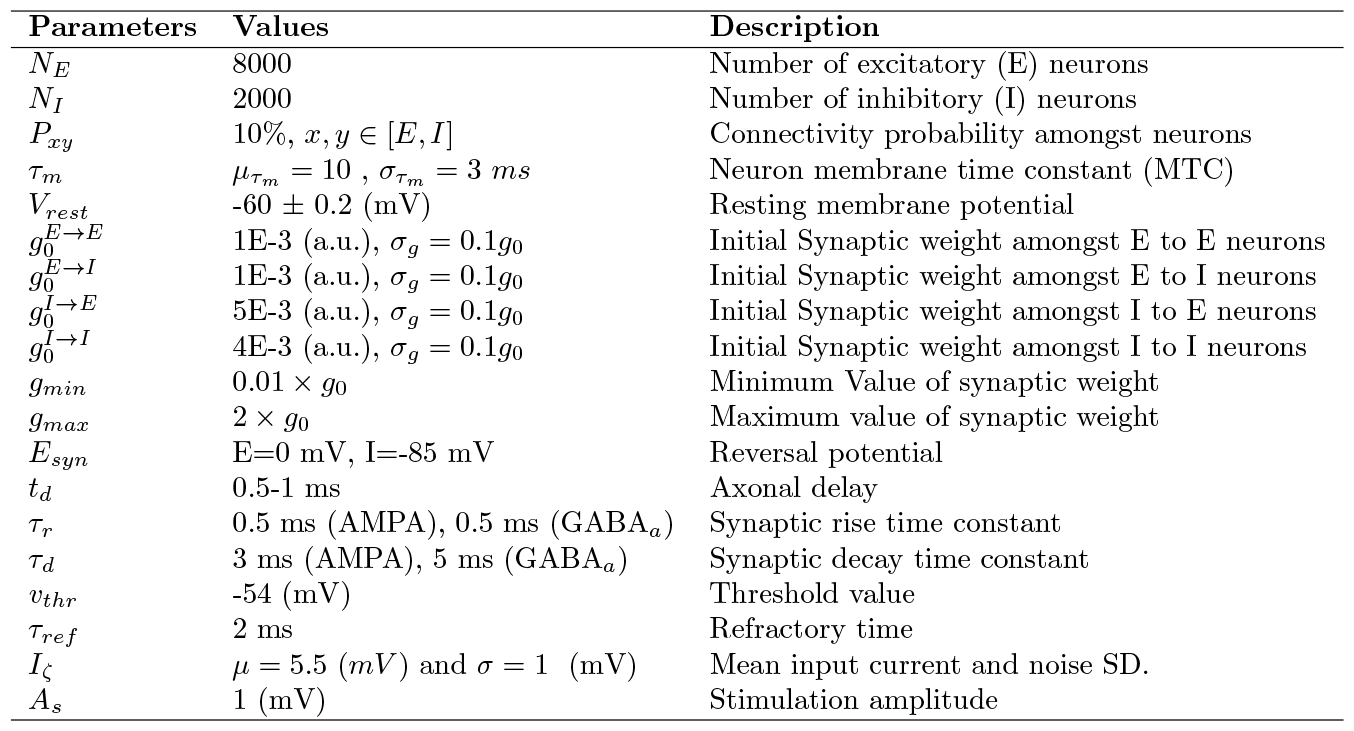
Parameters of the neuronal populations.

We subjected this network to periodic stimulation of various amplitude (*A*_*s*_) [49], and frequency (*ω*_*s*_), for a period of fifteen seconds (simulation time). We then compared changes between the dynamics observed before stimulation (i.e., pre-stimulation) and after stimulation (i.e., post-stimulation) over epochs of five seconds, respectively. To investigate the relationship between MTC heterogeneity and the persistence of stimulation-induced aftereffects, we plotted representative dynamics of the network in pre- (sham) and post-stimulation epochs in Fig. 1. We randomly selected 60 excitatory neurons, and compared both network connectivity and the relative magnitude of synaptic weights between pre-and post-stimulation epochs in Fig. 1(A1) and (B1), respectively. Comparing these panels, one can readily notice stimulation-induced changes in synaptic weights and/or connectivity persisting well after stimulation offset. This effect was found to be mediated by variability in MTC. Corresponding synaptic weights matrices are plotted in Fig. 1(A2) and (B2), respectively. Although most synaptic weight changes persisted after stimulation offset, synapses between neurons with high MTC mismatch were more persistent compared to their pre-stimulation values, as previously reported [18]. As shown in Fig. 1(A3) and (B3), the endogenous synchronous irregular activity present in the pre-stimulation period has been amplified in the post-stimulation epoch, accompanying a persistent increase in neuronal firing rates (Note that firing rates are lower than the network endogenous frequency as expected from irregular synchronous dynamics [50], i.e. the median of excitatory neurons firing rate is ∼ 0.5Hz for pre-stimulation and ∼1.5Hz for post-stimulation. see Fig. 1A4 and B4). The underlying population ‘s local field potential (LFP)(see Eq. 5 in *Materials and methods*) also exhibits a significant and persisting increase in spectral power, especially salient at the endogenous (i.e., resonant) oscillation frequency and outlasting stimulation duration (see Figs 1 A5 and B5, also A6 and B6). We generalized these results in Figs. S1, and S2, by choosing different distances to the threshold for each cell in the network, as well as introducing heterogeneity in the threshold values.

**Fig 1.**
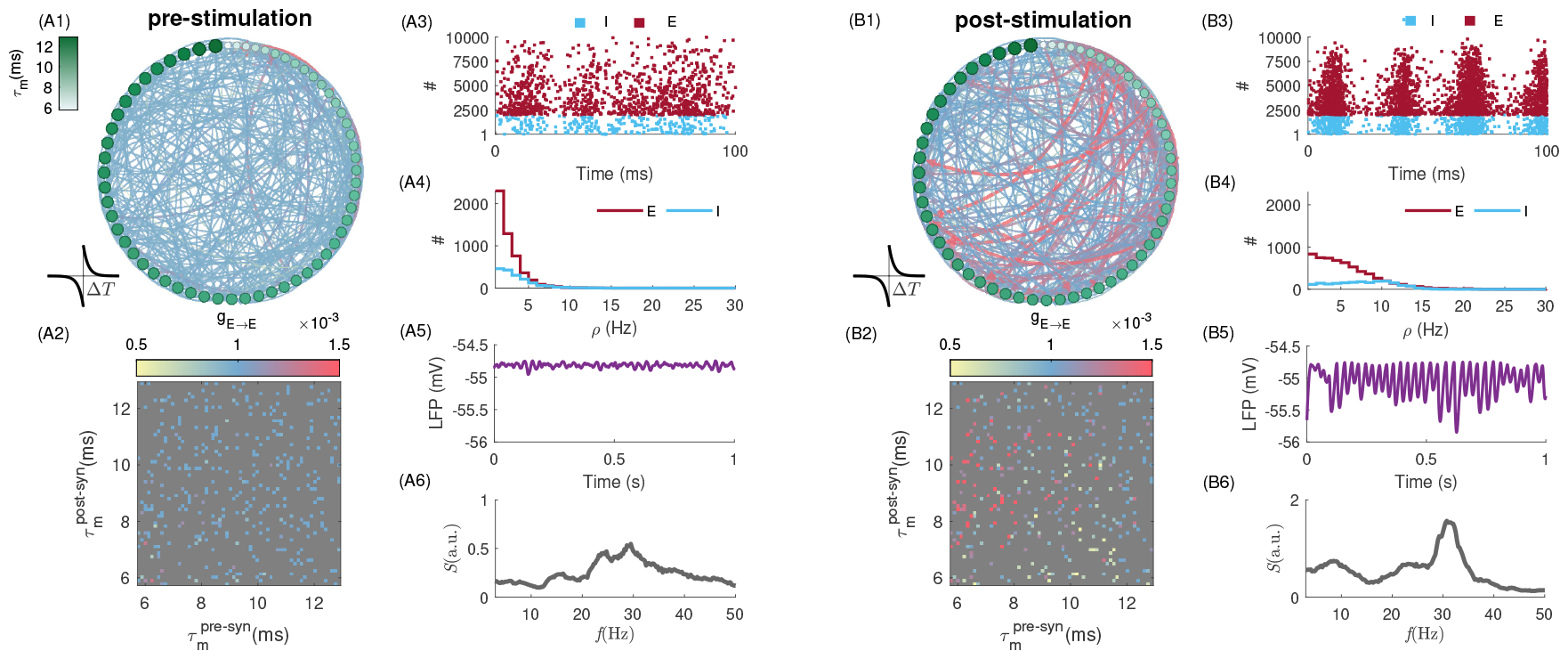
Amplified post-stimulation endogenous oscillations. (A1) and (B1) depict the pre- and post-stimulation population connectivity diagram, highlighting the changes in synaptic weights resulting from PBS. Here we plotted the connectivity amongst 60 randomly selected excitatory neurons during pre- (*t* = 5 *s*) and post-stimulation (*t* = 21 *s*) epochs, respectively. The neurons are sorted based on their MTC in a clockwise manner. The radius and colour of nodes indicated the change in the neuron ‘s MTC as the colorbar in (A1). The arrows indicate the connection from pre- to post-synaptic neurons. Synaptic weights are subjected to a Hebbian STDP (see 4; schematically shown as the inset plot). The arrows’ thickness and colour indicate the connection ‘s strength as colour-coded in (A2) and (B2), the corresponding synaptic weights matrices which are another representation of connectivity changes. The colorbar shows the strength of synaptic weights amongst pre-post neurons. (A3) and (B3) show the spiking activity of excitatory (E) and inhibitory (I) neurons in pre- and post-stimulation epochs, respectively. Note that the neurons’ spikes are plotted based on their MTC for each E (red dots) and I (blue dots) neuron, i.e., neurons with smaller MTCs have higher firing rates. (A4) and (B4) indicate neurons’ firing rates in the pre- and post-stimulation epoch, respectively. The population shows asynchronous irregular (AI) activity. Note that individual neuronal firing rates are smaller than the network endogenous oscillatory frequency. (A5) and (B5) show the LFP (see Eq. 5) for pre- and post-stimulation epochs, respectively. (A6) and (B6) show the resultant power spectrum of population activity in pre- and post-stimulation epochs, respectively. Here, *ω*_*s*_ = 25 *Hz*, and *A*_*s*_ = 1 *mV*. To plot the connectivity diagram (A1 and B1), we used freely available software *Gephi* [47].

Having identified post-stimulation amplification in endogenous oscillations, we next evaluated how this phenomenon depends on stimulation parameters. In Fig. 2 (A)-(B), we plot the peak LFP power for various stimulation frequencies, both during and after stimulation offset. Stimulating at frequencies ranging from *ω*_*s*_ = 1 *Hz* to *ω*_*s*_ = 40 *Hz* (*A*_*s*_ = 1 *mV*) invariably increases LFP power during entrainment, especially for stimulation frequencies near the resonant endogenous frequency. The effect carried over to the post-stimulation epoch: as can be seen in Fig. 2(B), peak power remained high around the population endogenous frequency despite no stimulation being present - indicative of stimulation-induced engagement of synaptic plasticity. Optimal post-stimulation peak power was observed at a stimulation frequency of *ω*_*s*_ ∼ 23 *Hz* which we note is different from the network endogenous oscillation observed before stimulation onset (*f* = 28 *Hz*). This indicates that stimulation-induced changes in synaptic coupling might be higher at non-resonant frequencies.

**Fig 2.**
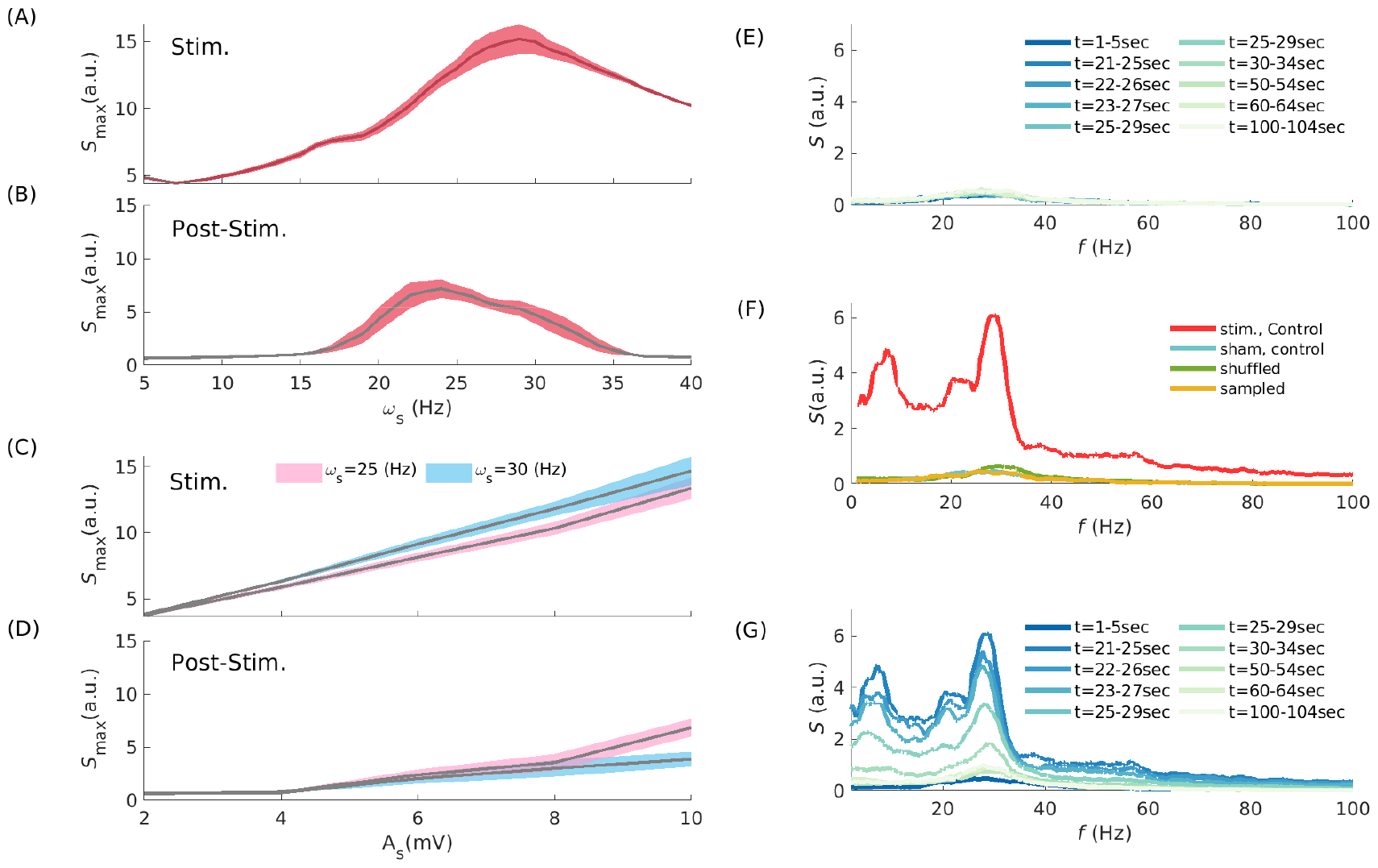
Effect of stimulation frequency and amplitude on post-stimulation activity. (A) and (B) show the maximum value of the LFP power spectrum at different stimulation frequencies during entrainment and post-stimulation epochs, respectively. Here *A*_*s*_ = 1 (*mV*). Note that the y-axes are in different scales. (C) and (D) show the maximum value of the LFP power spectrum while the amplitude of stimulation changes as the x-axis for entrainment and post-stimulation epochs, respectively, for *ω*_*s*_ = 25, and 30 *Hz*. (E), (F) and (G) display the power spectrum of LFP at different time points and situations. In (E), the stimulation is OFF, *ω*_*s*_ = 0 *Hz, A*_*s*_ = 0 (*mV*), and the figure shows the power spectrum of population oscillation within 2 minutes (simulation time) of free evolution. (F) Shuffling and re-sampling the synaptic weight matrix after turning off periodic stimulation suppresses spectral amplitude. (G) Illustrates the post-stimulation power changes observed at different time points. The colours, as the legend in (E), indicate the time intervals used to calculate the LFP power spectrum. In (E), (F), and (G), the stimulation was ON over *t* ∈ [5 20)*s* with *A*_*s*_ = 1 *mV* and *ω*_*s*_ = 25 *Hz*. The error bar, represented by the shaded area, denotes the standard deviation (SD) range around the trial-averaged values.

Stimulation amplitude is also crucial to elicit - and possibly maintain - persistent entrainment and associated changes in synaptic coupling. We plotted in Fig. 2(C) and (D) the peak LFP power as a function of stimulation amplitude (i.e., *A*_*s*_) both during and after stimulation offset. Two stimulation frequencies (i.e., *ω*_*s*_ = 25, and 30 *Hz*) were considered as they both reside within the range of frequencies for which the effect of post-stimulation LFP power is significant (see Fig. 2B). While peak LFP power increases linearly with stimulation amplitude during stimulation epochs (see Fig. 2C), a thresholding effect can be observed in the post-stimulation period. Indeed, a minimum stimulation amplitude appears to be required to cause post-stimulation LFP power amplification (Fig. 2D). These results indicate that high stimulation amplitude is required to modulate the neurons’ membrane potential, and spiking response, to cause changes in connectivity significant enough to yield persistent effects. The difference in LFP spectral power between the two selected stimulation frequencies (i.e., *ω*_*s*_ = 25 and 30 *Hz*) indicates that despite expected stimulation-induced resonance (here at *ω*_*s*_ = *f* = 30 *Hz*, see Fig. 2D), amplification may occur at different, non-resonant stimulation frequency. We however emphasize that stimulation-induced change in synaptic coupling may trigger shifts in endogenous oscillatory activity and vice versa.

We further investigated whether and how MTC heterogeneity is involved in generating those results. Is the persistent LFP power amplification observed post-stimulation due to a global, non-specific increase in synaptic coupling, or is it instead due to selective, MTC-mediated synaptic plasticity? To answer this question, we first explored the effects of STDP on post-stimulation power amplification. As shown in Fig. 2 (E), in the absence of stimulation (i.e., sham; *A*_*s*_ = 0) while the network remains exposed to STDP due to its own endogenous activity, no significant shift in LFP power can be observed, even after a prolonged learning period (three minutes, simulation time). This indicates that although synaptic coupling fluctuates endogenously over time, these do not result in any significant spectral fluctuations(cf. Fig. 1 B6).

Stimulation-induced amplification in post-stimulation power was found to rely heavily on selective synaptic modifications i.e. synapse-specific directional changes resulting from periodic entrainment of neurons possessing distinct MTCs [18]. To expose the role of such selectivity, we randomly shuffled synaptic weights amongst neurons of the same cell-type while preserving their overall statistics (see Materials and methods). Fig. 2 (F) compares the spectral power obtained without stimulation (sham control; *A*_*s*_ = 0 *mV*) and post-stimulation (stim control;*ω*_*s*_ = 25 *Hz, A*_*s*_ = 1 *mV*) conditions with those obtained by shuffling and/or sampling synaptic weights randomly while preserving their respective distributions, within and between cell types. To do this, we first calculated the synaptic weight distribution amongst all synaptic types (i.e., *E* → *E, E*→ *I*, and *I*→ *E*; *I*→ *I* remained unchanged). We next randomly shuffled synaptic weights in the network and examined whether post-stimulation oscillatory amplification could be observed over epochs of 4 seconds (no stimulation was applied during that period). As shown in Fig. 2 (F), no post-stimulation increase in power could be observed, indicating that while displaying the same overall statistics (i.e., same synaptic weight mean and variance), selective plasticity between neurons with distinct MTCs is essential in generating persistent entrainment. We pushed the analysis further and sampled synaptic weights independently, only using the cell-type specific distributions calculated above (i.e., agnostic of the actual values of those weights). The same result could be observed: in the absence of selectivity, post-stimulation oscillatory amplification vanishes.

Despite the significance of oscillatory amplification and its manifest reliance on MTC heterogeneity, all persistent after-effects reported were found to be transient [17, 24, 25] and dissipate over time after stimulation is turned off. Upon stimulation offset, prevailing endogenous synchronous irregular activity engages STDP to bring synaptic connectivity back to baseline [18] (see Fig. 2 G).

We examined the evolution of synaptic weights between all types of synapses in Fig. 3 and their distribution with respect to differences in MTCs, i.e., 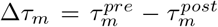. In Fig. 3, we plot synaptic weights for different stimulation frequencies i.e. *ω*_*s*_ =15 Hz (Fig. 3 A1-A4), 25 Hz (Fig. 3 B1-B4), and 35 Hz (Fig. 3 C1-C4). These frequencies were selected to help the comparison between the dynamics and resulting plasticity at stimulation frequencies that either amplify the post-stimulation power (i.e., *ω*_*s*_ = 25 *Hz*) and frequencies that do not (*ω*_*s*_ = 15, 35 *Hz*; see Fig. 2B). Although synaptic changes are noticeable in all of these cases, their relative magnitude and distribution were found to be highly frequency-specific. For instance, synaptic weights between excitatory and inhibitory neurons (i.e., *E* → *I*, (Fig. 3 A2, B2, C2) display a broader distribution of synaptic modifications at *ω*_*s*_ = 25 *H* (Fig. 3 B2) compared to other frequencies (Fig. 3 A2, C2). This indicates that the resonant stimulation frequency( ∼25 *Hz*), solicits MTC heterogeneity more strongly, leading to selective synaptic changes spanning a greater range of Δ*τ*_*m*_ and stronger power amplification (see Fig. 3 A4, B4, and C4). This is in contrast to Fig. 3 (A2), and (C2) where synaptic weight changes were more selective for negative Δ*τ*_*m*_. The same effect could be observed for synapses between different cell types: selective modification observed amongst *E*→ *E* and *I*→*E* synapses displayed a similar trend. In the last column, (A4), (B4), and (C4), for comparison purposes we plotted the pre- and post-stimulation power resulting from each stimulation frequency used (*ω*_*s*_ = 15, 35 *Hz*). Due to the aforementioned changes in synaptic weight, the strongest observed post-stimulation power occurred around non-resonant stimulation frequency (i.e., *ω*_*s*_ = 25 *Hz*; see Fig. 3 B4).

**Fig 3.**
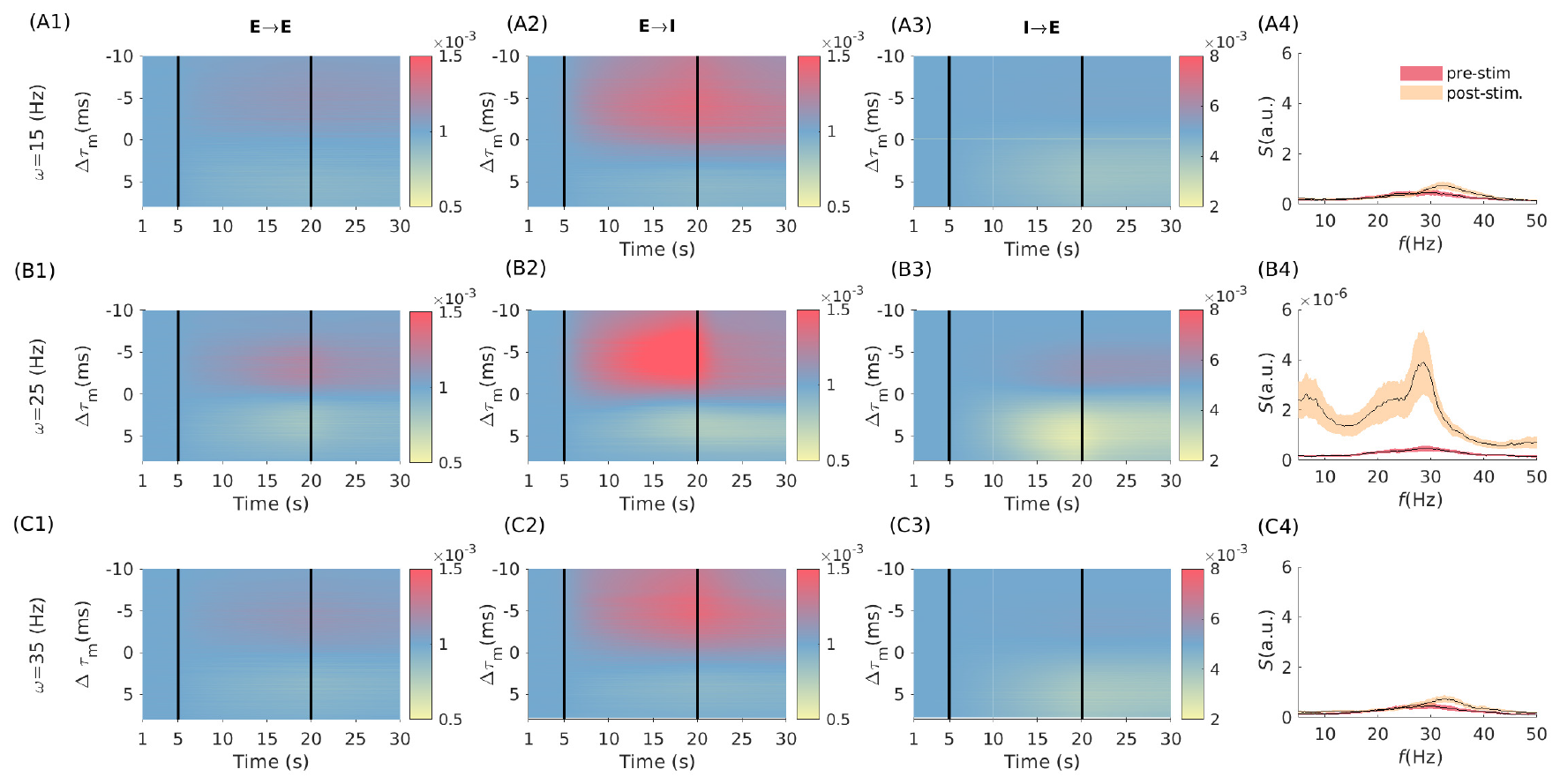
MTC heterogeneity shapes synaptic weights distribution and spectral power in a frequency- and cell-type specific manner. Figure groups A, B, and C (i.e., A1-A4) are related to the stimulation frequencies *ω*_*s*_ = 15, 25, and 35 *Hz*, respectively. The heat-map plots show the distribution of synaptic weights over time (x-axis) between synapses which we sorted according to their MTC difference (y-axis), 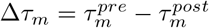. Figures in each of the columns (first, second, and third column), from left to right, depict the evolution of the synaptic weights between *E*→ *E, E*→ *I*, and *I*→ *E*, respectively, for the 30s. Vertical lines in each panel divided the simulation into three epochs: the pre-stimulation (*t* = [1 5)*s*), stimulation (*t* = [5 20)*s*), and post-stimulation (*t* = [20 30)*s*) epochs. In the last columns, (A4), (B4), and (C4), the power spectrum of neuronal population rhythm for pre- and post-stimulation epochs are plotted. For better comparison, we preserved the same y-axis range for all panels. The error bar, represented by the shaded area, denotes the standard deviation (SD) range around the trial-averaged

Heterogeneity amongst and between different cell types, either excitatory or inhibitory, have different consequences on the post-stimulation power. To quantify this, we explored in Fig. 4 the effects of cell-type MTC heterogeneity on post-stimulation LFP power. As shown in Fig. 4 (A), MTC heterogeneity among excitatory neurons enhances post-stimulation power, whereas increasing MTC heterogeneity among inhibitory neurons abolishes the effects (See A). The greater diversity observed among cortical interneurons compared to excitatory neurons [51], may hinder stimulation effects and possibly prevent power amplification. However, it should be noted that the frequency of stimulation is another factor that determines the stimulation effects. We measured this in Fig. 4 (B), where we varied the level of MTC heterogeneity of E and I neurons and the frequency of stimulation. A similar increase in MTC variability of E and I neurons contributes to the induction of post-stimulation effects over a wider stimulation frequency range. Having the same heterogeneity among inhibitory and excitatory neurons amplifies response power and therefore creates the necessary conditions for optimal synaptic weight changes, which ultimately leads to the amplification of oscillation power.

**Fig 4.**
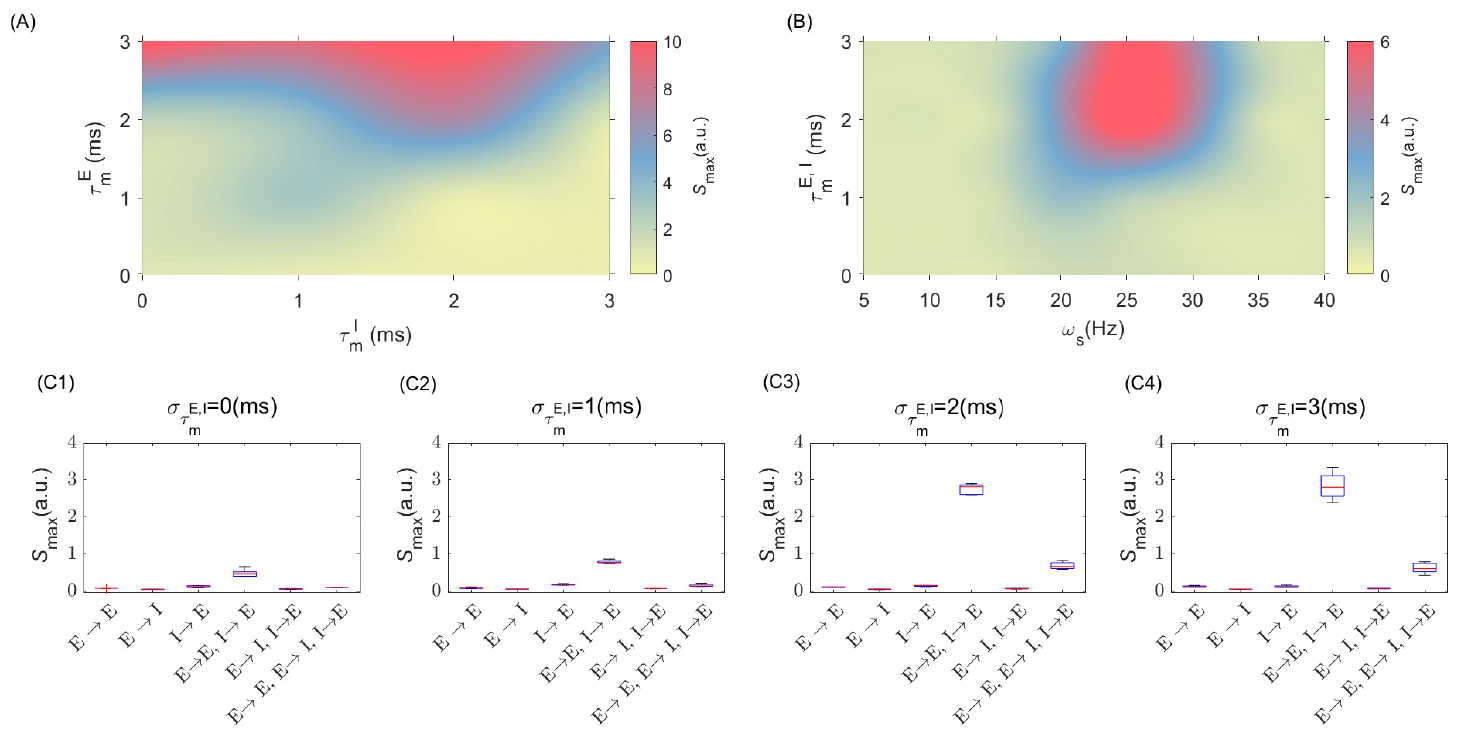
MTC heterogeneity amongst cell types modulates post-stimulation oscillation power. (A) shows the maximum power spectrum of the post-stimulation epoch when the MTC ‘s distribution for E and I neurons change as the x- and the y-axis, respectively. (B) show the maximum power spectrum of post-stimulation power when the stimulation frequency changes as the x-axis and the MTC ‘s distribution on E and I neurons changes as the y-axis. (C1) to (C4) show the changes in the maximum power spectrum of post-stimulation epochs when the STDP is just ON over indicated groups of neurons as x-axis at different values of 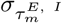, respectively.

Another way of examining these results is to consider plasticity amongst and between different cell types. To investigate which synapses are more significantly involved in mediating the post-stimulation aftereffects, we applied periodic electrical stimulation on the same population at different degrees of MTC heterogeneity while selectively turning ON and OFF STDP amongst different cell types. This enabled the identification of synapses whose plasticity is more significantly solicited during stimulation. In Fig. 4 (C1-C4), we show that plasticity amongst excitatory-to-excitatory (i.e., *E*→*E*) and between inhibitory-to-excitatory (i.e., *I* → *E*) neurons are more involved in the persistence of LFP power. The effect was also found to scale with the level of MTC heterogeneity across cell types (excitatory and inhibitory neurons) as of Fig. 4 (C1-C4) where the post-stimulation power amplified as we increased the 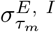. Introducing plasticity among inhibitory neurons, under the same conditions as those considered previously, is found to suppress the amplitude of post-stimulation aftereffects (see Fig. S3). The results suggest that blocking synaptic plasticity amongst specific synapses may lead to a significant increase in the post-stimulation power.

## Materials and methods

### Spiking neuron model

We modelled a population of excitatory and inhibitory Leaky-Integrate- and-Fire (LIF) neurons [45, 52]. The differential equation for the evolution of the membrane potential of each neuron is

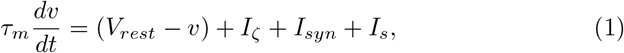

where *τ*_*m*_ is the MTC, *v* is the membrane potential, *V*_*rest*_ is the resting membrane potential, and *I*_*ζ*_ is the external current as white noise with mean value *μ* and standard deviation *σ. I*_*syn*_ represents the synaptic current. *I*_*s*_ is the stimulation-induced current, which is a sinusoidal input, i.e., *I*_*s*_ = *A*_*s*_ sin(2*πω*_*s*_*t*), where *A*_*s*_ and *ω*_*s*_ are the amplitude of the periodic signal, and the angular frequency respectively. When a neuron crosses the threshold value *v*_*thr*_ = −54 *mV*, it spikes and its membrane potential resets to resting value *V*_*rest*_ = −60 *mV* and remains there for *τ*_*ref*_ = 2 *ms* representing the neuronal refractory period. Although having larger refractory periods alters the neurons’ firing, the results remain consistent (not shown). The parameters are in the physiological range [53] and summarized in table 1. Changing the distance to threshold (i.e., |*v*_*rest*_ − *v*_*thr*_|) and/or introducing the heterogeneity in neurons’ spiking threshold value (i.e., *v*_*thr*_), did not qualitatively alter these results, as shown in Figs. S1 and S2, respectively. The total simulation time, unless otherwise stated, is 30 seconds, including pre-stimulation (Sham epoch): *t* ∈[0 5)*s*, stimulation epoch: *t*∈[5 20)*s*, and post-stimulation epoch: *t*∈ [20 30]*s*. The total synaptic current for neuron *i* is given by

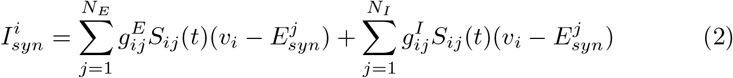

where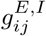 are synaptic weights matrix associated with connections between either excitatory (E) and inhibitory (I) pre-synaptic neurons towards a post-synaptic neuron *i*. The sum is taken over *N*_*E*_ excitatory and *N*_*I*_ inhibitory pre-synaptic neurons. The reversal potential, *E*_*syn*_, for E and I neurons are 0 (*mV*) and −80 (*mV*), respectively. The synaptic response function *S*_*ij*_(*t*) for connections from neuron *j* to neuron *i* is modeled as

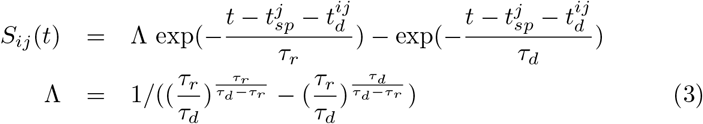

where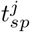is the spiking time of *j*^*th*^ neuron, and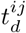 is the axonal delay between pre-synaptic neuron, *j*, and post-synaptic neuron, *i*. The *τ*_*r*_ and *τ*_*d*_, are rise and decay synaptic time constants, respectively, associated with GABA_*a*_ and AMPA receptors (see table 1) [53].

The synaptic weights matrix is randomly chosen from a normal distribution with mean and standard deviation as given in table 1 (sham control). In *shuffled*, first we let the simulation run for 20*s*, and instantaneously shuffled the synaptic weights at the beginning of the post-stimulation epoch. We shuffled synaptic weights within and between cell types. In the *sampled* case, we took the following procedure: We let the population in *stim. controlled* evolve for 20s (5s pre-stimulation, and 15s stimulation epochs). We then calculated the distribution of the synaptic weights at the end of the stimulation epoch, computing its mean and standard deviation. Then we used these values to randomly sample synaptic weights within each synapse category (*E* → *E, E* → *I*, and *I* → *E*) using this normal distribution.

### Spike Timing-Dependent Plasticity (STDP)

Plasticity in our population amongst connected neurons is modelled using Hebbian spike-timing dependent plasticity [53, 54]. To avoid biased synaptic changes (i.e., preferential LTP/LTD), we chose a symmetric STDP Hebbian learning rule [46, 55] with soft bands [56]. The synaptic weights dynamic in our model follows the below equations [18]

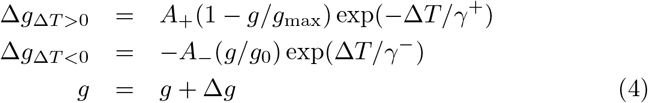

The *γ*^+^ and *γ*^*−*^ are STDP decay time constants. 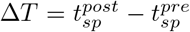 is the time difference between the spiking time of post- and pre-synaptic neurons. Whenever Δ*T* is positive (negative), the synaptic weight between *pre* to *post* neurons gets potentiated (depressed). The constant *g*_max_ denotes the maximum achievable synaptic weight, while *g*_0_ denotes the initial synaptic weight, taken from narrow Gaussian distribution across all synaptic connections before learning (see table 1).

Synaptic changes are bounded within a physiologically relevant range by setting the maximum and minimum value of synaptic weight to *g*_*min*_ = 0.01*g*_0_ and *g*_*max*_ = 2*g*_0_. Therefore, every time the synaptic weights over-pass these limits, this condition will impose the value to those mentioned limits to ensure the synaptic weight remains between boundaries. Baseline synaptic coupling and threshold were selected to set the network in a weak coupling, sub-threshold regime, in which an isolated pre-synaptic spike does not guarantee post-synaptic firing. Despite weak synaptic coupling, the afferent synchronous synaptic input each neuron receives from the rest of the network is comparable to the stimulation amplitude, i.e. the average of maximum synaptic input and its standard deviation is ∼ 0.5 *mV* and ∼ 0.1 *mV*, respectively. Throughout this report, we used Eq. 4 for synaptic modification, and our choice of STDP parameters are *A*_+_ = 2*A*_*−*_ = 0.02 and *γ*_*±*_ = 10 *ms*.

### Network model

We modelled a randomly connected sparse network of 10000, leaky-integrate-and-fire (LIF) neurons (see Eq. 1) with a 4:1 ratio of E (8000) and I (2000) neurons with a fixed connection probability of 0.1 [50, 57, 58]. To balance physiological relevance and computational tractability for the network sizes we used the LIF neurons model [59]. The synaptic weights and other neurons’ parameters have been selected within the reported physiological range [58] to be in line with previous studies on LIF cortical network models (see [53, 59–61] and references therein), and are further summarized in table 1. To study the effect of MTC heterogeneity, we randomly sampled neuronal MTCs (*τ*_*m*_) from Gaussian distribution with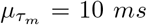, and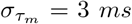 unless otherwise specified. The resulting population exhibits a synchronous irregular activity (SI).

### Power spectral analysis

To perform spectral analysis of the network mean activity, we first calculated the local field potential (LFP),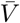 as the weighted ensemble average of the membrane potential i.e.,

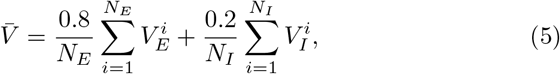

where the relative proportion of excitatory (0.8) versus inhibitory(0.2) cells is taken into consideration. The power spectral density of 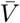 was averaged over 10 independent trials in which the same stimulation protocol is applied, but using different baseline connectivity, synaptic weights, and noise realizations. For the purpose of Figs. 1, 2, and 4, we further took the average of the power spectral density with a moving average window (with MATLAB *smooth* function) with *σ* = 1.5 *Hz*, that provided us with a smoothed power-frequency curves (for instance, see Fig. 2 B6).

## Discussion

To better understand the mechanism underlying persistent post-stimulation amplification in oscillatory activity observed in experiments [15,62], we extended the framework of selective STDP [18] in a synchronous, sparsely connected neuronal network of heterogeneous spiking neurons. We computationally showed that in the presence of endogenous synchronous activity, near-resonant periodic stimulation may amplify post-stimulation power through selective synaptic changes, whose magnitude and direction rely on intrinsic differences in MTC. Stimulation at the near-resonant frequency was found to engage STDP so that the population expresses higher endogenous oscillatory power (see Fig. 2 B), resulting in transient yet prolonged overlasting effects. We confirmed that selective, directional changes in synaptic coupling - both within and between cell-types - are responsible for such amplification, while any shuffled, randomly assigned synaptic weights, or intrinsic synaptic weight changes in the absence of stimulation, are insufficient for generating aftereffects on their own(see Fig. 2 E, F, and G). The level of heterogeneity in neuronal MTC was found to determine the efficacy of stimulation on post-stimulation power magnitude and duration (see Fig. 4 A). Indeed, in a homogeneous network (i.e., where the MTCs are identical), neurons respond similarly to a given stimulus. Because of the symmetric nature of our STDP rule [53, 54, 56], such homogeneity might prevent stimulation-induced synaptic plasticity, even in the presence of noise. This means that a minimum level of heterogeneity is essential for pushing STDP in one direction or another, especially while interacting with time-varying inputs. Taken together, these results reveal one potential mechanism behind the effectiveness of PBS for therapeutic purposes, specifically the stabilization of stimulation effects on neural dynamics and connectivity. We argue that heterogeneity in neuronal time scales represents a dominant contributor mediating PBS efficacy, affirming the neurophysiological bases of persistent entrainment towards the development and/or optimization of clinical interventions.

It should be noted that the results we report here extend to a broad range of endogenous frequencies. For instance, networks expressing oscillations within the alpha range may need different stimulation frequencies to solicit selectivity in synaptic plasticity [36]. Interestingly, while stimulating at resonant/endogenous frequency expectedly yields higher entrainment [36] (see Fig. 2 A), this does not always accompany significant post-stimulation aftereffects (see Fig. 2 B). We point out that our simulations also support a state-dependent dependence on stimulation efficacy. Indeed, weak background synaptic activity resulted in a high signal-to-noise ratio i.e. stimulation-induced modulation in neuronal membrane potential was significant enough to trigger depolarization and hence recruit STDP. In the presence of strong synaptic activity, however, the effects may fade away [30, 36]. We also emphasize that to engage populations expressing a wide range of MTC, stimulation amplitude must scale accordingly, potentially influencing neuronal firing rates [18]. The precise relationship between stimulation frequency, synaptic plasticity, and persistent entrainment remains to be fully explored.

Nonetheless, our model suffers from limitations. First, we considered a neuronal network with random local connectivity, among cell types (*E*→*E, E*→ *I, I* →*E*, and *I*→*I*). The more realistic network as observed experimentally [63] has a different connectivity distribution which should be considered in later investigations. Note that the connectivity distribution, either locally or globally, could lead to different axonal delay distributions among neurons which then may influence the synaptic plasticity dynamics [64]. Second, the symmetric Hebbian STDP rule, and our assumption that all synapses obey the same rule, are limiting the generality of our results. Future investigations need to consider the large variety of synaptic plasticity mechanisms between cell types [54, 65] and the possible heterogeneity in STDP parameters. Note that introducing both Hebbian and Anti-Hebbian plasticity for efferent inhibitory synapses (i.e. *I* → *I* and *I*→*E*) [54, 65–67] yields qualitatively similar results, yet their amplitude is suppressed (see Fig. S3). These results showcase the importance of synaptic dynamics on the emergence and persistence of oscillatory activity in recurrent neural networks and warrant further investigation.

Synaptic plasticity selectivity is not limited to heterogeneity in MTC: other sources of heterogeneity such as the resting membrane potential, rheobase, and/or spiking threshold, may promote cell-to-cell differences in spike timing. Lastly, we have mapped neuron ‘s MTCs using a normal distribution, whose variance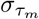 (i.e., scaling with the degree of heterogeneity) alters the number of synapses that can be effectively modified by stimulation. However, similar to natural phenomena, the MTC distribution may be better fitted using a gamma or lognormal distribution [18, 68, 69].

Another limitation arises from our choice of using the same MTC distribution for both excitatory and inhibitory neurons. This choice was motivated by the need to balance physiological relevance and computational tractability - as well as limiting the dimensionality of the analysis. While the introduction of cell-type specific MTC distributions would certainly influence our results, we note that by construction, excitatory and inhibitory cells in our network already display differences in firing rates (e.g., see Fig. 1). Further investigations are warranted to thoroughly examine such additional sources of heterogeneity. We however hypothesize that as long as the overall activity of the neuronal population remains within an oscillatory synchronous irregular state, characterized by a low level of coherency, similar results would be observed.

## Conclusion

Brain stimulation techniques offer non-invasive treatments for brain-related disorders. The promising results in the application of these techniques attracted a wide range of interdisciplinary researchers to investigate the response of brain cells to these interventions and devise more effective and reliable methods. Towards this goal, our study expanded the knowledge of how periodic stimulation may enhance and stabilize post-stimulation effects. Our results emphasize the importance of neural timescale variability in the interaction between synaptic plasticity and PBS. Overall, our results elucidate one potential mechanism by which PBS affects neural population connectivity, and conditions under which such intervention can lead to persistent overlasting effects.

## Acknowledgments

We thank the National Research Council of Canada (NSERC GRANT RGPIN-2017-06662) as well as the Canadian Institute for Health Research (CIHR GRANT NO PJT-156164)(AH) for funding. The funders had no role in study design, data collection and analysis, decision to publish, or preparation of the manuscript.

## Code accessibility

The codes are available publicly on GitHub https://github.com/arefpz/neuronal_population.

---

**Fig S1.**
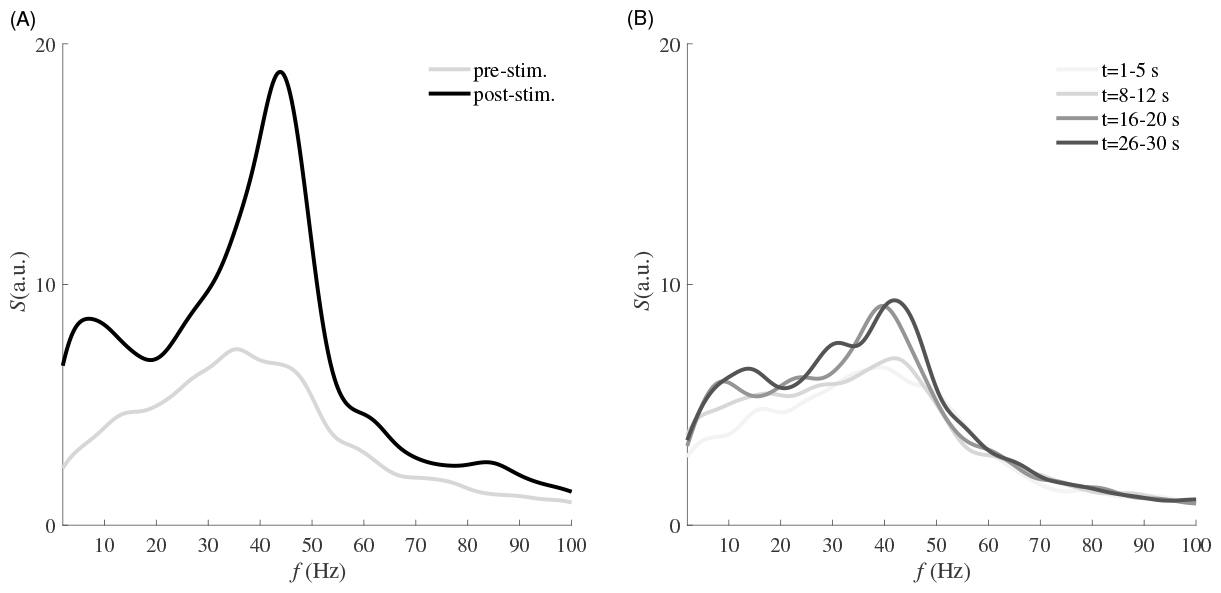
Larger distance to threshold values has no significant impact on the pre-post oscillation amplitude. The population oscillation frequency in pre and post-stimulation epochs is shown as grey and black lines, respectively, when the distance to the threshold has increased to 16 (*mV*). To push the neurons toward the spiking threshold region, we changed the input current in Eq. 1 to 15.5 (*mV*). In this case, the ratio of stimulation amplitude and distance to the threshold is reduced to ∼ 0.06 (originally was ∼ 0.17).

**Fig S2.**
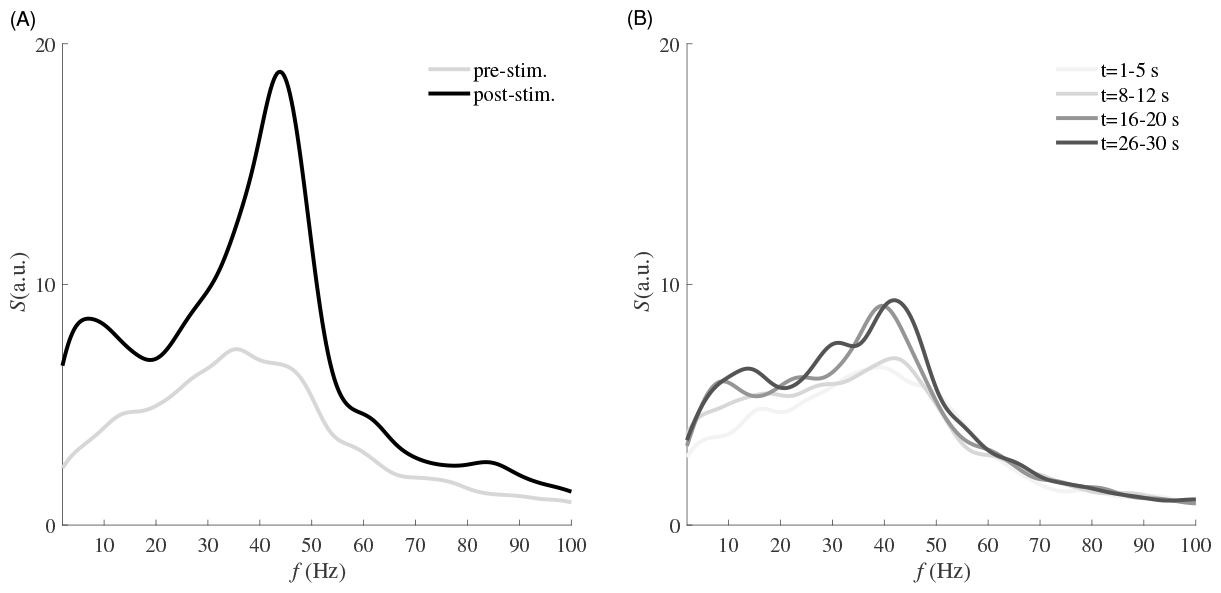
Effect of heterogeneity in neuron ‘s spiking thresholds. By introducing spiking threshold heterogeneity in both excitatory and inhibitory cells, the population oscillates at a different endogenous frequency (*f* = 43 *Hz*). Applying stimulation at the peak frequency, (i.e., *ω*_*s*_ ≈*f* = 43 *Hz*) for 15 *s* enhanced the oscillation amplitude in (A). (B) In the presence of threshold heterogeneity and STDP, but in the absence of stimulation, the population oscillations reach a steady state at the same endogenous frequency (i.e., *f* = 43 *Hz*) and amplitude. The threshold values were sampled from a normal distribution with *μ*_*thr*_ = −54 (*mV*), *σ*_*thr*_ = 2 (*mV*).

**Fig S3.**
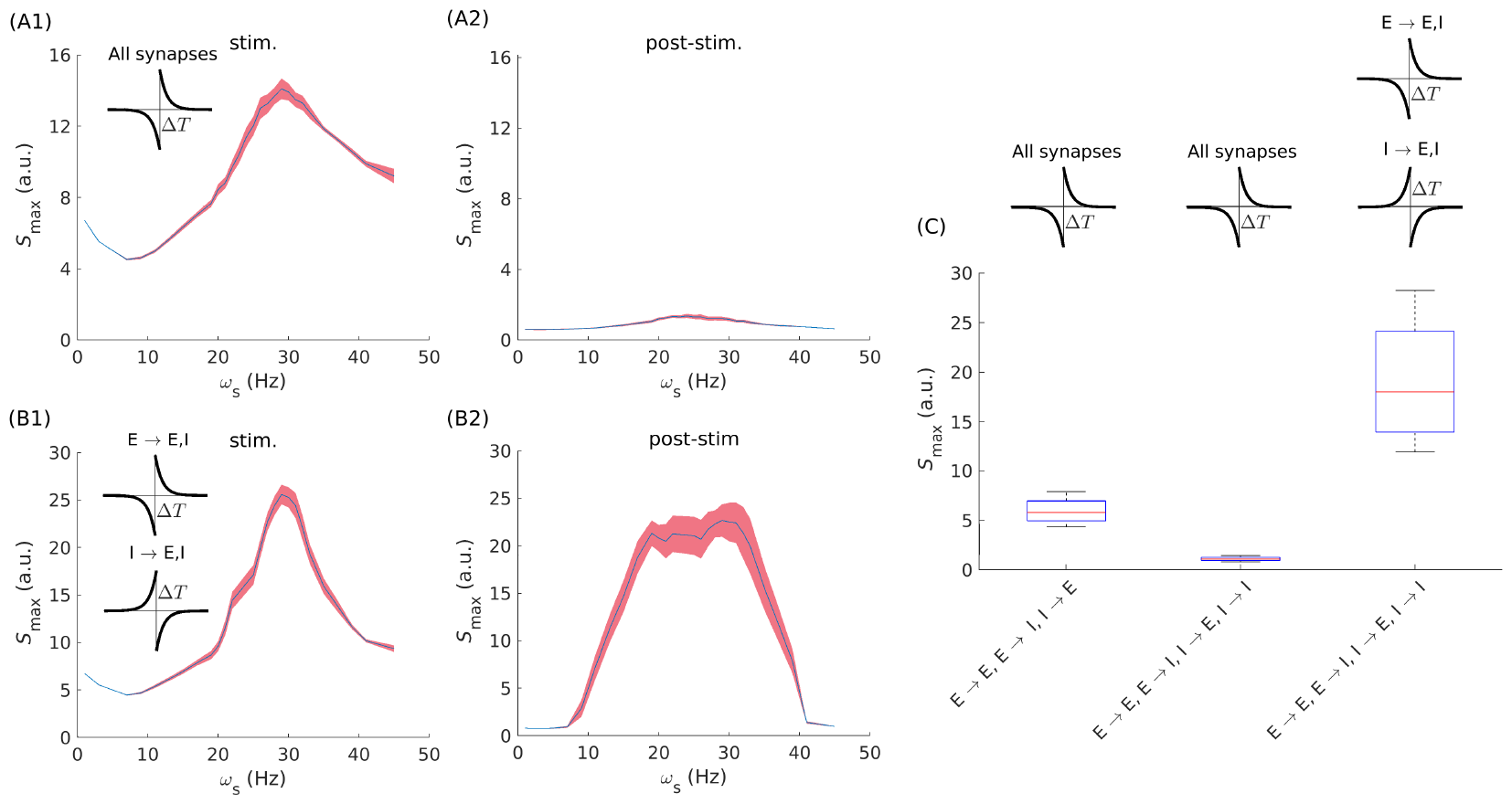
Hebbian and anti-Hebbian STDP effects on post-stimulation power spectrum. (A1) and (A2) show the power spectrum of stimulation and post-stimulation epochs when the synaptic weights are modified with the Hebbian STDP rule as shown inset plot in (A1). (B1) and (B2) show the power spectrum of stimulation and post-stimulation epochs, when the Hebbian STDP rule is active for efferent synapses from excitatory neurons, and anti-Hebbian rule for efferent synapses from inhibitory neurons as depicted in inset plots in (B1). (C) Comparison of the post-simulation power spectrum for three cases as noted in the x-axis. The results showcase the effects of synaptic plasticity on post-stimulation power modification. Introducing the plasticity among inhibitory neurons reduced the post-stimulation power amplitude in comparison to Fig. 2 where the *I* →*I* plasticity was absent. The STDP type (Hebbian, anti-Hebbian) not only affects the post-stimulation epoch power amplitude but also affects the entrainment power amplitude (see the A1 and B1), which needs further investigation for clarification.

