## Supplementary Figures for "Amplifying post-stimulation oscillatory dynamics by engaging synaptic plasticity with periodic stimulation: a modelling study": supplementstuff.tex

\documentclass{article}

\usepackage{graphicx}
%\usepackage{xr}

%\usepackage{subcaption}
\begin{document}
% \section*{Supplementary Figures}
\begin{figure*}[!h]
    \renewcommand{\thefigure}{S1}
    \centering
    \includegraphics[width=12cm]{s1.eps}
    \caption{{\bf Larger distance to threshold values has no significant impact on the pre-post oscillation amplitude.} The population oscillation frequency in pre and post-stimulation epochs is shown as grey and black lines, respectively, when the distance to the threshold has increased to $16~(mV)$. To push the neurons toward the spiking threshold region, we changed the input current in Eq. 1 to $15.5~(mV)$. In this case, the ratio of stimulation amplitude and distance to the threshold is reduced to $\sim 0.06$ (originally was $\sim 0.17$).}
    \label{fig:S1_fig}
\end{figure*}
\begin{figure*}[!h]
    \renewcommand{\thefigure}{S2}
    \centering
    \hspace*{-2cm}
    \includegraphics[width=12cm]{s2.eps}
    \caption{{\bf Effect of heterogeneity in neuron's spiking thresholds.} By introducing spiking threshold heterogeneity in both excitatory and inhibitory cells, the population oscillates at a different endogenous frequency (${\it f}=43~Hz$). Applying stimulation at the peak frequency, (i.e., $\omega_s \approx {\it f} = 43~Hz$) for $15~s$ enhanced the oscillation amplitude in (A). (B) In the presence of threshold heterogeneity and STDP, but in the absence of stimulation, the population oscillations reach a steady state at the same endogenous frequency (i.e., ${\it f}=43~Hz$) and amplitude. The threshold values were sampled from a normal distribution with $\mu_{thr}=-54~(mV),~\sigma_{thr}=2~(mV)$.}
    \label{fig:S2_fig}
\end{figure*}
\begin{figure*}[!h]
    \renewcommand{\thefigure}{S3}
    \centering
    \hspace*{-2cm}
    \includegraphics[width=14cm]{s3.jpg}
    \caption{{\bf Hebbian and anti-Hebbian STDP effects on post-stimulation power spectrum.} (A1) and (A2) show the power spectrum of stimulation and post-stimulation epochs when the synaptic weights are modified with the Hebbian STDP rule as shown inset plot in (A1). (B1) and (B2) show the power spectrum of stimulation and post-stimulation epochs, when the Hebbian STDP rule is active for efferent synapses from excitatory neurons, and anti-Hebbian rule for efferent synapses from inhibitory neurons as depicted in inset plots in (B1). (C) Comparison of the post-simulation power spectrum for three cases as noted in the x-axis. The results showcase the effects of synaptic plasticity on post-stimulation power modification. Introducing the plasticity among inhibitory neurons reduced the post-stimulation power amplitude in comparison to Fig. 2 where the $I \rightarrow I$ plasticity was absent. The STDP type (Hebbian, anti-Hebbian) not only affects the post-stimulation epoch power amplitude but also affects the entrainment power amplitude (see the A1 and B1), which needs further investigation for clarification.}
    \label{fig:S3_fig}
\end{figure*}

\end{document}
