## Supplementary figures and images for "Amplifying post-stimulation oscillatory dynamics by engaging synaptic plasticity with periodic stimulation: a modelling study"

### s3.jpg

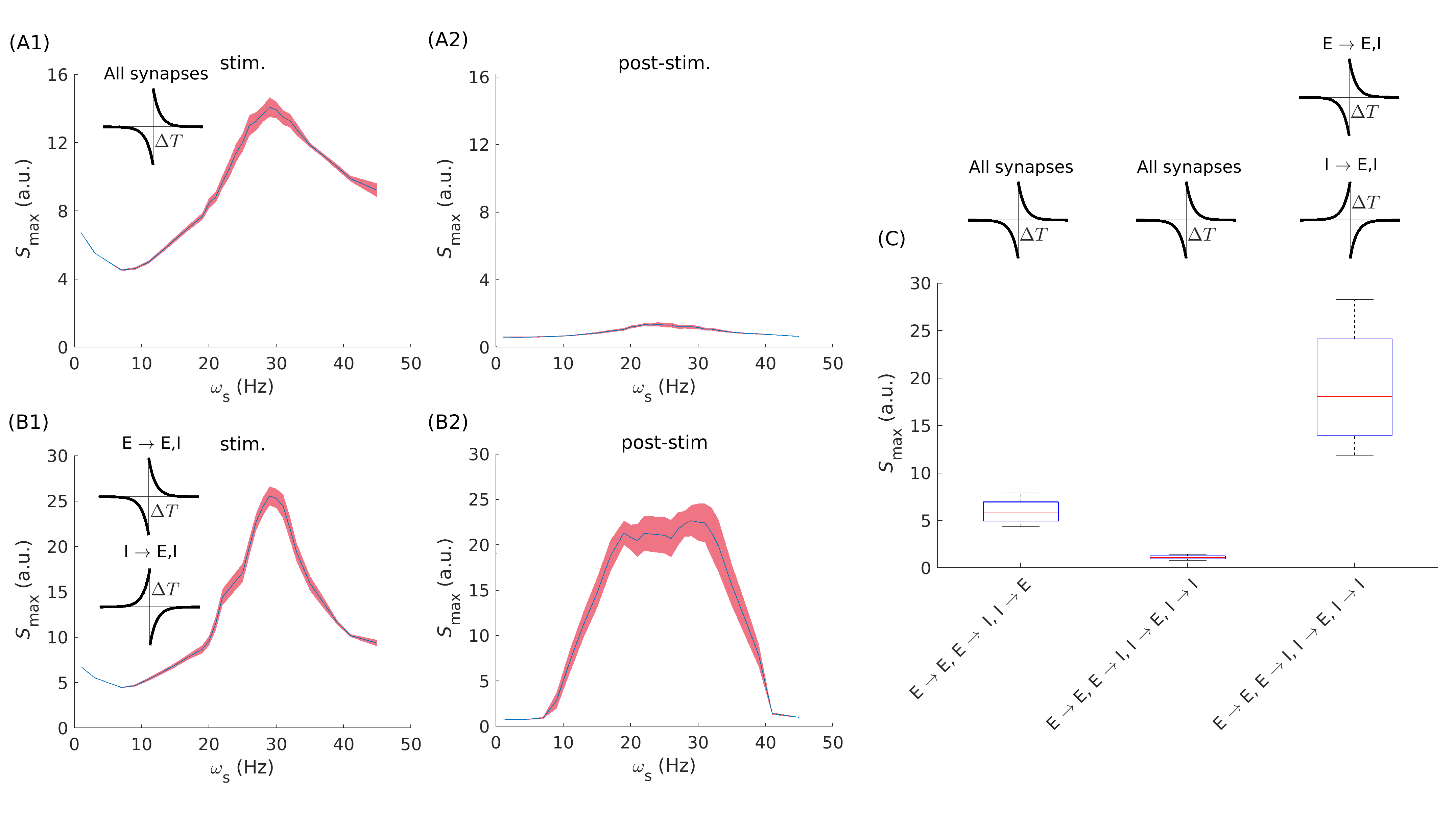
